# Splice junction-centric approach to identify translated noncanonical isoforms in the human proteome

**DOI:** 10.1101/372995

**Authors:** Edward Lau, Yu Han, Damon R. Williams, Rajani Shrestha, Joseph C. Wu, Maggie P. Y. Lam

## Abstract

RNA sequencing has led to the discovery of many transcript isoforms created by alternative splicing, but the translational status and functional significance of most alternative splicing events remain unknown. Here we applied a splice junction-centric approach to survey the landscape of protein alternative isoform expression in the human proteome. We focused on alternative splice events where pairs of splice junctions corresponding to included and excluded exons with appreciable read counts are translated together into selective protein sequence databases. Using this approach, we constructed tissue-specific FASTA databases from ENCODE RNA sequencing data, then reanalyzed splice junction peptides in existing mass spectrometry datasets across 10 human tissues (heart, lung, liver, pancreas, ovary, testis, colon, prostate, adrenal gland, and esophagus) as well as generated data on human induced pluripotent stem cell directed cardiac differentiation. Our analysis identified 1,108 non-canonical isoforms from human tissues, including 253 novel splice junction peptides in 212 genes that are not documented in the comprehensive Uniprot TrEMBL or Ensembl RefSeq databases. On a proteome scale, non-canonical isoforms differ from canonical sequences preferentially at sequences with heightened protein disorder, suggesting a functional consequence of alternative splicing on the proteome is the regulation of intrinsically disordered regions. We further observed examples where isoform-specific regions intersect with important cardiac protein phosphorylation sites as well as generated data on human induced pluripotent stem cell directed cardiac differentiation. Our results reveal previously unidentified protein isoforms and may avail efforts to elucidate the functions of splicing events and expand the pool of observable biomarkers in profiling studies.

## Introduction

The human proteome contains many more variant isoforms than there are coding genes (Aebersold et al., 2018; Smith and Kelleher, 2018). Alternative splicing is a mechanism through which a single gene can create multiple transcripts, and changes in alternative splicing has been widely implicated in development, aging, and diseases (van den Hoogenhof et al., 2016; Lee and Rio, 2015). RNA sequencing data have uncovered over 100,000 alternatively spliced transcripts from virtually all multi-exonic genes in the human genome (Pan et al., 2008; Wang et al., 2008). Yet relatively few alternative transcripts have been detected as protein product. Mass spectrometry is the preeminent tool for unbiased protein identification, but faces technical challenges in identifying alternative isoforms. Some factors that have impeded isoform identification include their low abundance (Blencowe, 2017) and frequent sequence incompatibility with proteases (Wang et al., 2018b). Arguably most importantly however, shotgun proteomics relies on protein sequence databases for protein identification, hence a continued lack of precise isoform sequences precludes isoform identification by search algorithms. As a result of these technical challenges, the protein molecular functions of most alternative splicing events remain uncharacterized, and our field lacks a systematic view on how alternative splicing can rewire the proteome functions (Tress et al., 2017b, 2017a).

There have been several approaches proposed to improve mass spectrometry identification of protein isoforms, including the curation of splice variant databases (Mo et al., 2008; Tavares et al., 2014) and de novo six-frame translation of genomic sequences (Fermin et al., 2006; Power et al., 2009). More recently, RNA sequencing has been leveraged with some successes to create sample-specific peptide databases representing expressed transcripts including splicing variants (Cifani et al., 2018; Sheynkman et al., 2013; Zickmann and Renard, 2015). Initial work showed that this approach can identify novel peptides not found in annotated sequence databases (Ning and Nesvizhskii, 2010), hinting at its potential utility for discovering protein isoforms. Thus far however, the majority of studies of this type have largely been performed in transformed human cell lines or cancerous tissues (Evans et al., 2012; Koch et al., 2014; Ning and Nesvizhskii, 2010; Sheynkman et al., 2013) which are known to express aberrant splice variants. Moreover, many custom transcript-guided databases remain imprecise and contain large proportions of indiscriminately translated sequences that likely cannot be found in the biological sample (e.g., translations from multiple frames), suggesting there is a need for continued refinement of *in silico* translation and evaluation methods.

Here we apply an approach that focuses on translatable splice junction pairs from RNA sequencing data to guide protein isoform identification. By focusing on alternative splicing events with appreciable read counts rather than individually assembled transcripts, we restrict the inflation of the resulting isoform databases, which induce false positives in database search (Alfaro et al., 2014; Ning and Nesvizhskii, 2010). We utilized the generated FASTA databases to recover alternative protein isoforms across diverse existing mass spectrometry data sets on healthy non-cancerous human tissues as well as a generated dataset on human induced pluripotent stem cell directed cardiac differentiation. The results suggest that this approach is able to extract both annotated non-canonical protein isoforms as well as uncharacterized novel junction peptides from mass spectrometry experiments including from previously unidentified spectra.

## Results

### Generation of junction-centric protein sequence databases

We applied a custom computational workflow to translate alternative splice junctions to protein sequences in silico (Figure 1). Differential exon inclusion analysis is a frequently used tool in transcriptomics to assess the percent-spliced-in (PSI) of splice events and differential exon usages across samples. We reasoned that by focusing our analysis around alternative junction pairs rather than all assembled transcripts, we can target relevant alternative splicing events that are likely to be appreciably variant within a tissue and create more precise sequence databases than indiscriminate in silico translation. We retrieved ENCODE RNA sequencing data on the GTEx tissue collection of human heart, lungs, liver, pancreas, transverse colon, ovary, testis, prostate, and adrenal gland, each containing 101-nt paired-end total RNA sequencing data from two biological replicate donors that passed ENCODE consortium-wide quality control. Sequencing reads are mapped to a reference human genome using STAR to identify transcript reads spanning splice junctions. To collect alternative splice events, we use rMATS to gather and count junction-specific reads for splicing events including alternative 3’ splice site (A3SS), alternative 5’ splice site (A5SS), mutually exclusive exons (MXE), skipped exon (SE), and retained intron (RI). rMATS as well as similar tools such as MISO are commonly employed to identify and quantify alternative splice events using GTF-annotated exons or directly from RNA reads, and return the sequencing read counts across alternative junction pairs (exon-included junction and exon-excluded junction) in each splicing events (Figure 1A).

**Figure 1.**
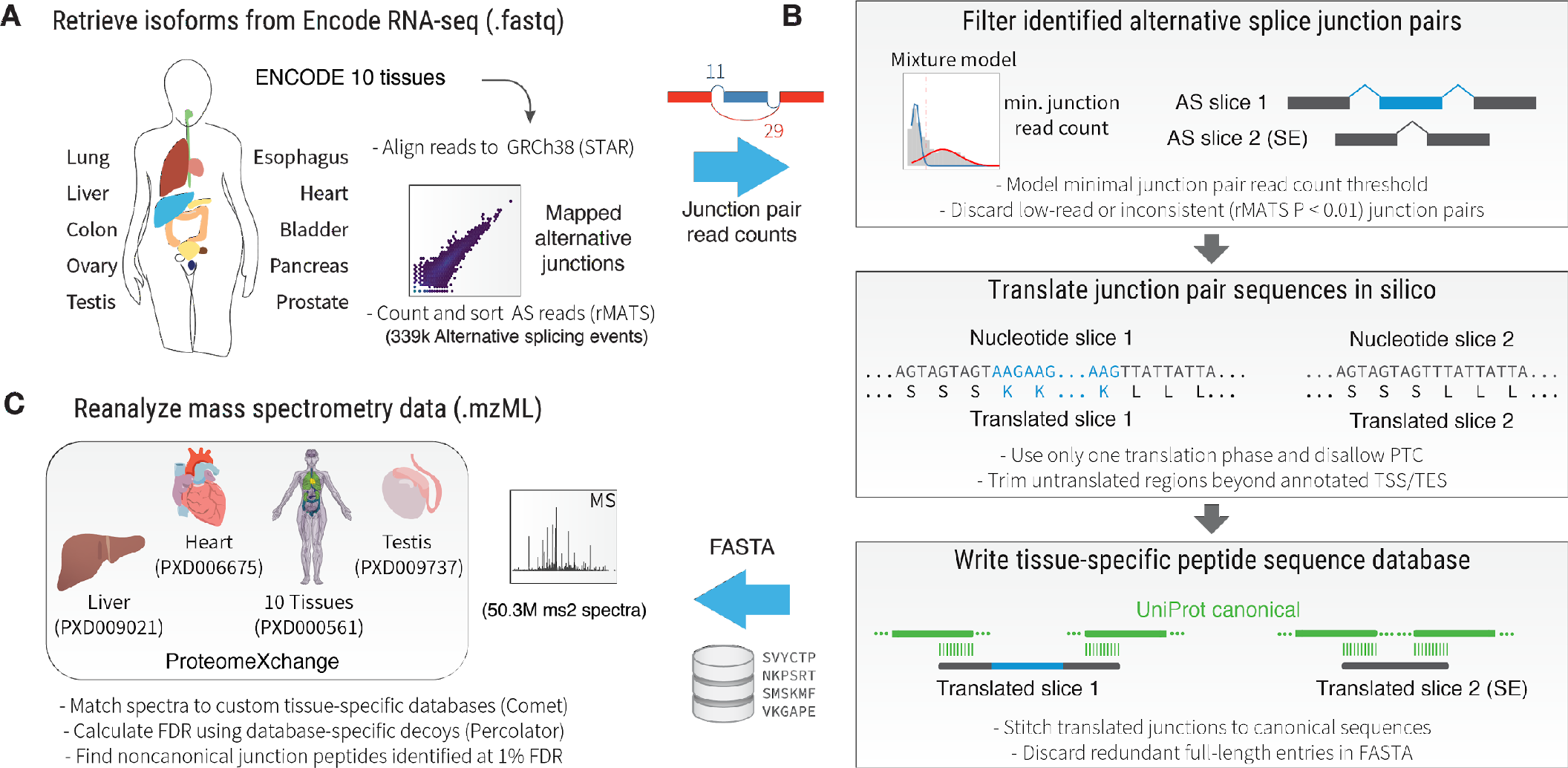
Splice-junction centric approach to identify protein isoforms. Schematic of the described approach. **A.** ENCODE RNA sequencing data across human tissues are mapped to GRCh38. Alternative splicing events and their corresponding junction read counts are extracted. **B.** Alternative splicing pairs are filtered for junction read counts and consistency. Candidate junctions were trimmed using Ensembl GTF-annotated translation start sites (TSS) and translation end sites (TES), then translated in-frame using either GTF-annotated reading frames or by choosing the translation frame that does not encounter premature termination codons (PTC). The translated junction pairs are extended to encompass the full length of the protein. **C.** The created tissue-specific FASTA databases containing splice-junction based isoform entries are used to reanalyze non-canonical protein isoforms across existing and newly generated datasets.

To restrict database sizes, we implemented four criteria to select splice junction pairs that are more likely detectable in mass spectrometry experiments (Figure 1B). (i) The skipped junction read counts of an alternative splice event must pass a sample-specific threshold. (ii) We use the statistical model implemented in rMATS to remove events with significantly different exon usage across technical and biological replicates in the same healthy tissue (P ≤ 0.01). (iii) We prioritize transcripts with known annotated translation start sites and frame and could be translated in-frame without premature termination codons (PTC). Where an unambiguous annotated translation frame is not available, we use one frame that results in the longest translatable sequence with no PTCs. (iv) To ensure that reliable junction peptides can be identified that span constitutive and alternative exons, we require both translated slices in a splice pair (each containing one upstream exon, the alternative exons, and one downstream exon) to be joined end-to-end back to full length canonical sequences from UniProt/SwissProt canonical sequences through a 10-amino-acid overhang. Orphan splices that are not stitchable back to canonical sequences are discarded, and redundant sequences are combined. Junction pairs that are translated in pair and extended using canonical SwissProt sequences to cover the entire protein sequence are written in a FASTA file for database search (Figure 1C).

From the ENCODE RNA sequencing data, we mapped 73,111 alternative splicing events per tissue on average, with fewest in the adrenal gland (66,160) and most in the testis (91,895). The most common type of alternative splicing events was SE, accounting for 65.1% of all identified events, followed by A3SS (10.8%), then RI (10.1), MXE (7.3%) and A5SS (6.7%). The mapped splice junctions show a broad distribution of skipped junction read counts (Figure 2A). As cellular transcription is noisy and previous work has established that mass spectrometry-based proteomics primarily detects the gene products form a population of more highly expressed transcripts (Ramakrishnan et al., 2009), we determined the optimal junction read count threshold to minimize inclusion of non-translatable junctions from encumbering database size and inflating false positive. Indeed we observed that the database entry counts scale with read count filter in a log-linear relationship in the analyzed RNA sequencing data (Figure 2B). We therefore removed low abundance junctions based on the excluded junction read count of the two alternative splice junctions created by a splice events, such that only alternative splice events where both alternatives may be expressed at likely detectable levels are retained. To identify the optional read count thresholds, a Gaussian mixture model grouped splice junctions by log read counts into groups corresponding to low and highly expressed isoform junctions, and a cutoff was applied that corresponded to a posterior probability of 0.95 of a splice junction falling into the highly-expressed group (Figure 2C), which in the ENCODE heart RNA sequencing-generated FASTA database corresponded to alternative splice events where both alternative junctions had at least 4 mapped reads.

**Figure 2.**
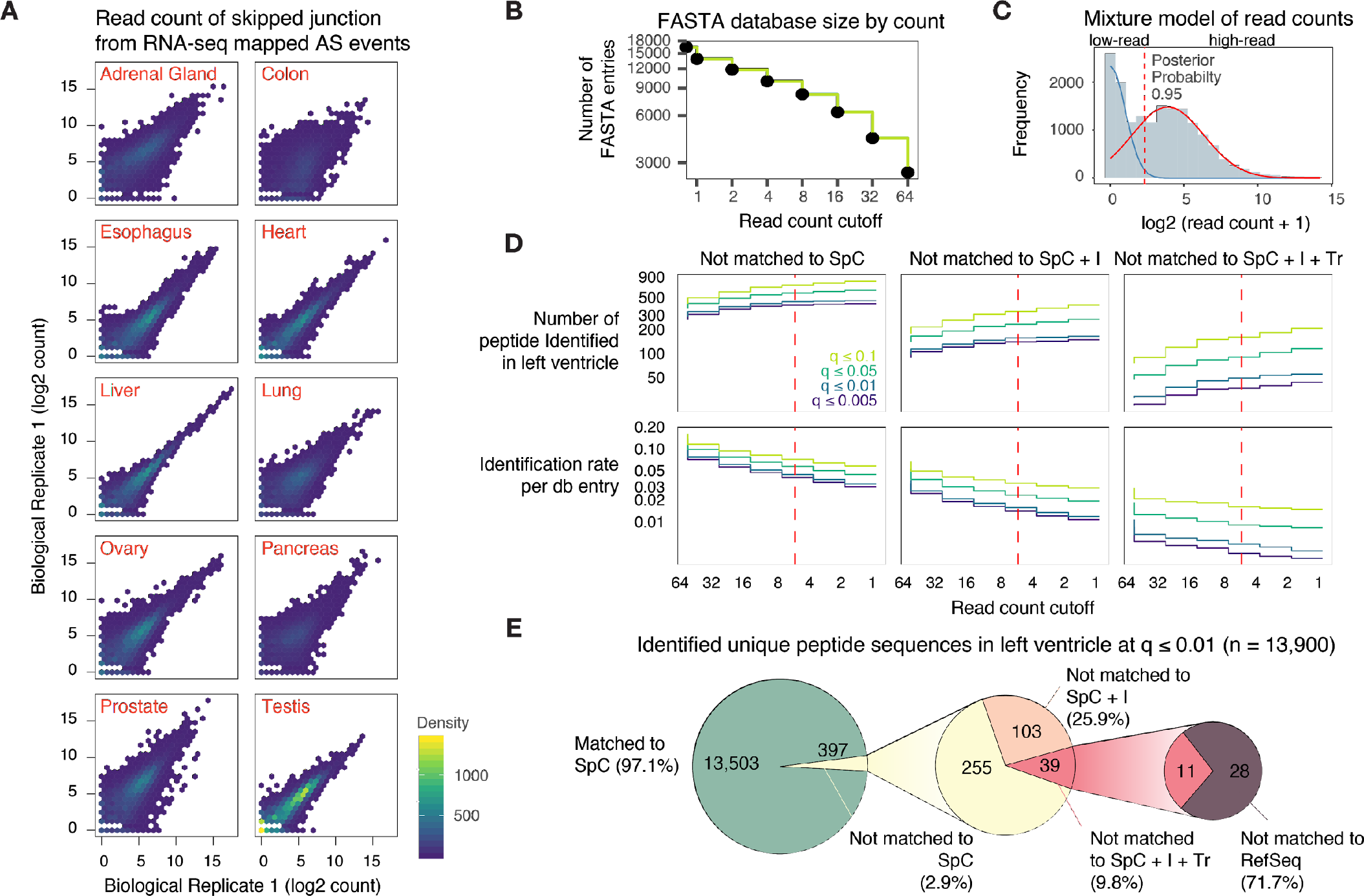
Junction filtering produces size-restricted FASTA databases. **A.** Density scatter plot showing the distribution of log2 read counts in the skipped junction read counts of all identified alternative splicing events in replicate RNA-seq data. **B.** Relationship between number of FASTA entry and minimal skipped junction read count threshold following in silico translation of the ENCODE human heart data set. Inclusion of low-read junctions increases database sizes. **C.** Gaussian mixture fitting overlaid on the skipped junction read counts of all alternative splicing events in the human heart database. Dotted line denotes the threshold chosen. **D.** Number of identified non-canonical isoform peptide sequences in a pilot reanalysis of a mass spectrometry dataset on the human heart left ventricle under different junction read count threshold. Color denotes Percolator FDR cutoff calculated with the aid of a database-specific reverse decoy. **E.** Proportion of identified distinct peptide sequences in the left ventricle dataset (13,900 total) not matchable to SwissProt canonical (SpC), SwissProt canonical and isoform (SpC + I), TrEMBL (Tr), or RefSeq.

We evaluated how the read count filter modified the number of identifiable splice junction peptides in a pilot reanalysis of human heart left ventricle mass spectrometry data using the custom heart isoform database, focusing on junction peptides that match to non-canonical isoforms not found in the commonly used curated sequence database UniProt SwissProt *Homo sapiens* canonical. We saw that the number of identifiable junction peptides gradually plateaued out at 4 count cutoff at more stringent significance cutoff (Percolator q ≤ 0.01) but continued to rise in lower-confidence matches (q ≥ 0.05), indicating an inflation of false positives when low-read junctions are included. At the same time, the number of sequences identified per FASTA entry continued to fall with expanded database sizes, hence the cutoff chosen struck a trade-off between identification and false positive rates (Figure 2D). In total, the search identified 13,900 distinct peptide sequences at 1% false discovery rate (FDR) using a reverse decoy database followed by filtering by Percolator, 397 of which (2.9%) were not matched to SwissProt canonical sequences (Figure 2E). Out of those sequences, 142 (35.7%) not matched to the curated SwissProt canonical + isoform database and 39 (9.8%) were not matched the automatically annotated larger sequence collection TrEMBL. Taken together, these results suggest that the approach is able to identify non-canonical and novel isoform sequences at low FDR.

### Identification of non-canonical splice junctions across tissues

We proceeded to build custom isoform databases for the other analyzed tissues. The filtering strategy reduced the number of entries in the resulting sequence database (Figure 3A). For instance, the human heart-specific database contains a representative 13,816 entries following filtering. For comparison, UniProt Swissprot sequence database catalogs 42,237 protein entries in the human reference proteome from 20,226 coding genes (20,225 canonical + 22,012 isoform sequences). The TrEMBL component of the UniProt database, containing un-reviewed sequences, houses additionally 93,555 sequences and Ensembl RefSeq comprises 113,620 sequences. Other large collections exist from automatic annotation of deposited sequences, but it is unclear whether the majority of sequences are bona fide isoforms, fragments, polymorphisms, or redundant entries. Across all tissues, the databases contain on average 11,911 splice-junction specific protein sequence entries, with the human pancreas-specific database containing the fewest protein sequences (6,309) and the testis containing the most entries (19,285). Hence all generated databases here are markedly sparser than SwissProt, TrEMBL and RefSeq, and indiscriminate three-frame unstranded translation of transcripts. This is expected because each tissue is expected to only express a subset of genes in the human genome due to tissue-specific epigenetic landscape and gene regulation, and RNA sequencing data provide information only on the specific genes that are expressed above threshold.

**Figure 3.**
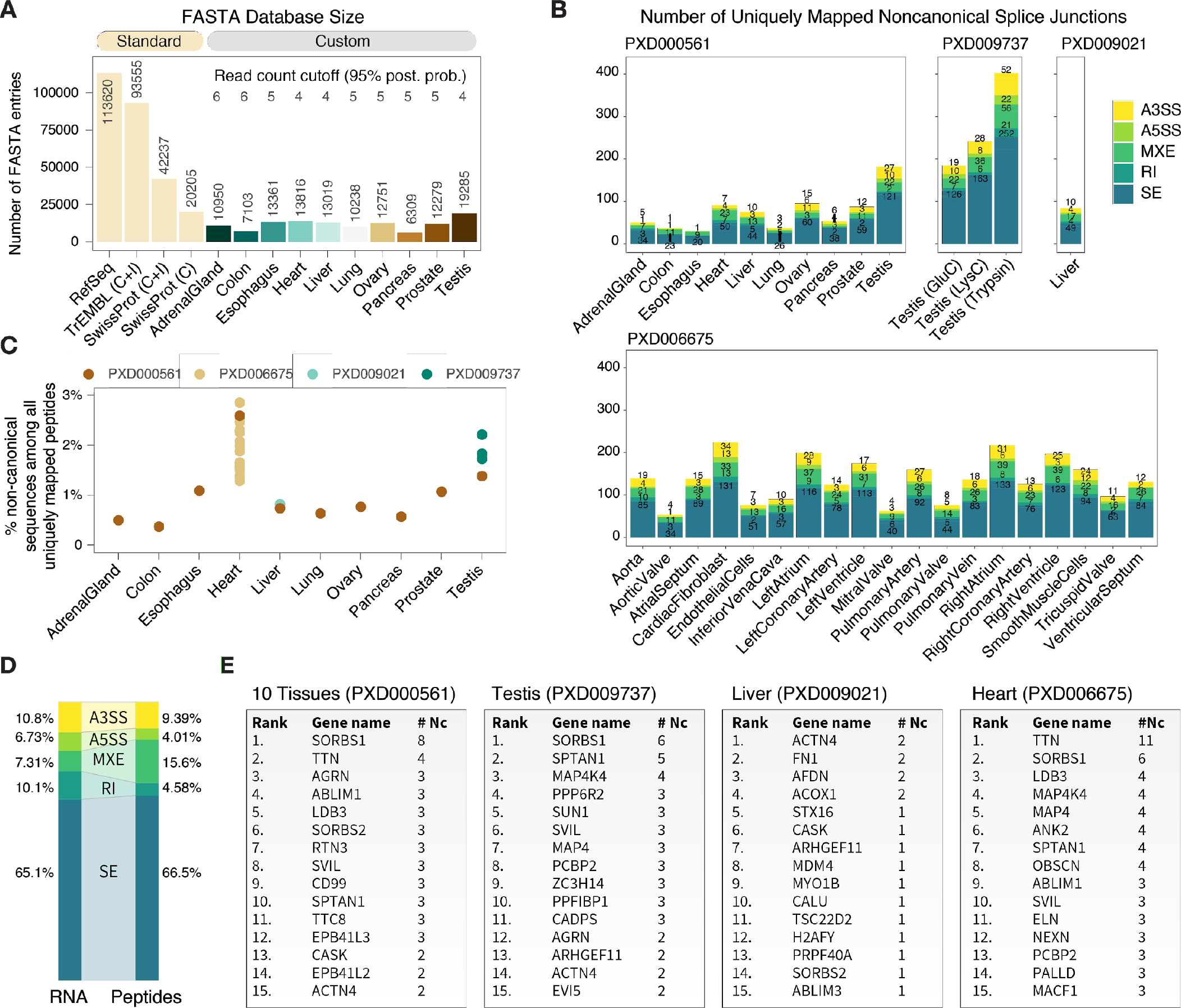
Non-canonical protein isoforms in the human proteome. **A.** Comparison on the number of entries in curated standard databases (RefSeq TrEMBL, SwissProt canonical + isoform and SwissProt canonical) vs. the custom generated tissue-specific databases. All generated databases are smaller than SwissProt. **B.** The number of uniquely-identified non-canonical junction peptides at 1% FDR across tissues in PXD000561, PXD009737, PXD009021, and PXD006675, including non-canonical sequences corresponding to known and documented on SwissProt isoforms as well as novel sequences. Color corresponds to alternative splicing type (A3SS, A5SS, MXE, RI, and SE); most identified non-canonical isoform peptides corresponded to skipped exon alternative splicing events. **C.** Proportion of distinct peptides uniquely mappable to non-canonical isoforms per tissue. The heart and the testis appear to be particularly enriched in non-canonical isoforms. Color corresponds to datasets. **D.** Proportion of alternative splicing types in RNA sequencing data (left) compared to identified non-canonical peptides (right), showing higher translatable rate for mutually exclusive exons (MXE). **E**. Tables showing the top 15 genes associated with the most identified non-canonical isoforms in each of the analyzed datasets.

We used the custom databases to perform a secondary analysis on four mass spectrometry datasets containing high-resolution Orbitrap FT/FT spectra on normal human tissues, comprising one dataset on 10 matching human tissues (PXD000561) (Kim et al., 2014), one on human testis using three proteases (PXD009737) (Sun et al., 2018), one on human liver using extensive fractionation (PXD009021), and one on the human heart dissected into 16 anatomical regions and 3 isolated cell types (PXD006675) (Doll et al., 2017). In total, we reprocessed 1,284 mass spectrometry experiment files with 50.3 million fragmentation spectra. In the heart which was the tissue with the most comprehensive reanalyzed data, 5,731 of 6,351 (90.4%) genes translated with at least one isoform in the heart database were mappable to an identified peptide in at least one isoform, whereas up to 23% of all translated isoforms are uniquely identifiable by a unique junction peptide or splice-specific peptide.

Because we translated both junction sequences in an alternative splice event in pairs, unique mappable peptides represent isoform-specific peptides and splice junctions. In total, we identified 2307 distinct and uniquely mappable peptides at 1% FDR which were not found in SwissProt canonical data, belonging to up to 1,108 non-canonical isoforms in 870 genes. (Figure 3B) In eight of the analyzed tissues, approximately 1% of distinct peptides that were mappable to a unique database entry correspond to a non-canonical isoform, whereas this proportion is markedly higher in the testis and the heart, suggesting alternative splicing may play pronounced roles in the proteomes of these two tissues (Figure 3C). Most of identified non-canonical peptides (66.5%) arose through skipped exon events, in line 65.1% of all alternative splicing events being skipped exons in the RNA sequencing data; mutually exclusive exons appeared to have higher translational potential (15.3% peptides vs. 7.3% at the RNA level), whereas retained introns produced relatively few translation products (4.6% non-canonical peptides vs. 10.1% in at the RNA level) (Figure 3D). We found a number of proteins with multiple identifiable non-canonical proteins including SORBS1 and MAP4K4 across multiple tissues, but unique multi-isoform proteins also exist including OBSCN and TTN in the heart (Figure 3E). The protein with the most isoforms identified is titin, which is also the largest protein encoded in the human genome with the most exons, whose splicing has been widely implicated in congenital heart diseases (Guo et al., 2012). Proteins with multiple non-canonical isoforms are found in diverse pathways including muscle contraction, metabolism, and signaling.

### Novel identified sequences in existing mass spectrometry data

Intriguingly, we also find splice junction peptide sequences that are novel to comprehensive databases. Among the non-canonical isoform sequences, we found 914 peptides identified at 1% FDR that are not matched to any entry in either SwissProt canonical or isoform sequences, belonging to 454 genes. Out of these, 338 peptides are also not matched to any entry in the TrEMBL isoform and canonical sequence database, which encompasses all SwissProt entries plus computationally annotated and unreviewed sequences. This result nominate a subset of TrEMBL sequences as likely uncharacterized splice isoforms, and provides tentative evidence for the protein-level existence of hundreds of isoform sequences not documented in UniProtKB. Among the TrEMBL-novel peptides, 253 belonging to 212 genes are further not matched to the larger Ensembl RefSeq collection. These novel peptides are found in every tissue and sub-anatomical regions analyzed with particular enrichment in the testis, which is known to differ more markedly from other tissues in alternative splicing pattern (Figure 4A). We note that the peptides for novel sequences across tissues had higher adjusted P values and posterior error probability than other non-canonical peptides that are found in TrEMBL, although a portion of novel peptides with high confidence are also clearly observable (Figure 4B). A lower score distribution for novel peptides may be due to their lower abundance or the known enrichment of lysines at splice junctions producing miscleavages (Wang et al., 2018b), which are penalized by the Percolator algorithm (The et al., 2016). Regardless of search engine scores, assignment of variant peptides demands caution and consideration of confounding causes including mass shifts due to post-translational modifications or single amino acid variant polymorphisms. Hence to further evaluate the novel peptide matches, we considered several additional lines of evidence.

**Figure 4.**
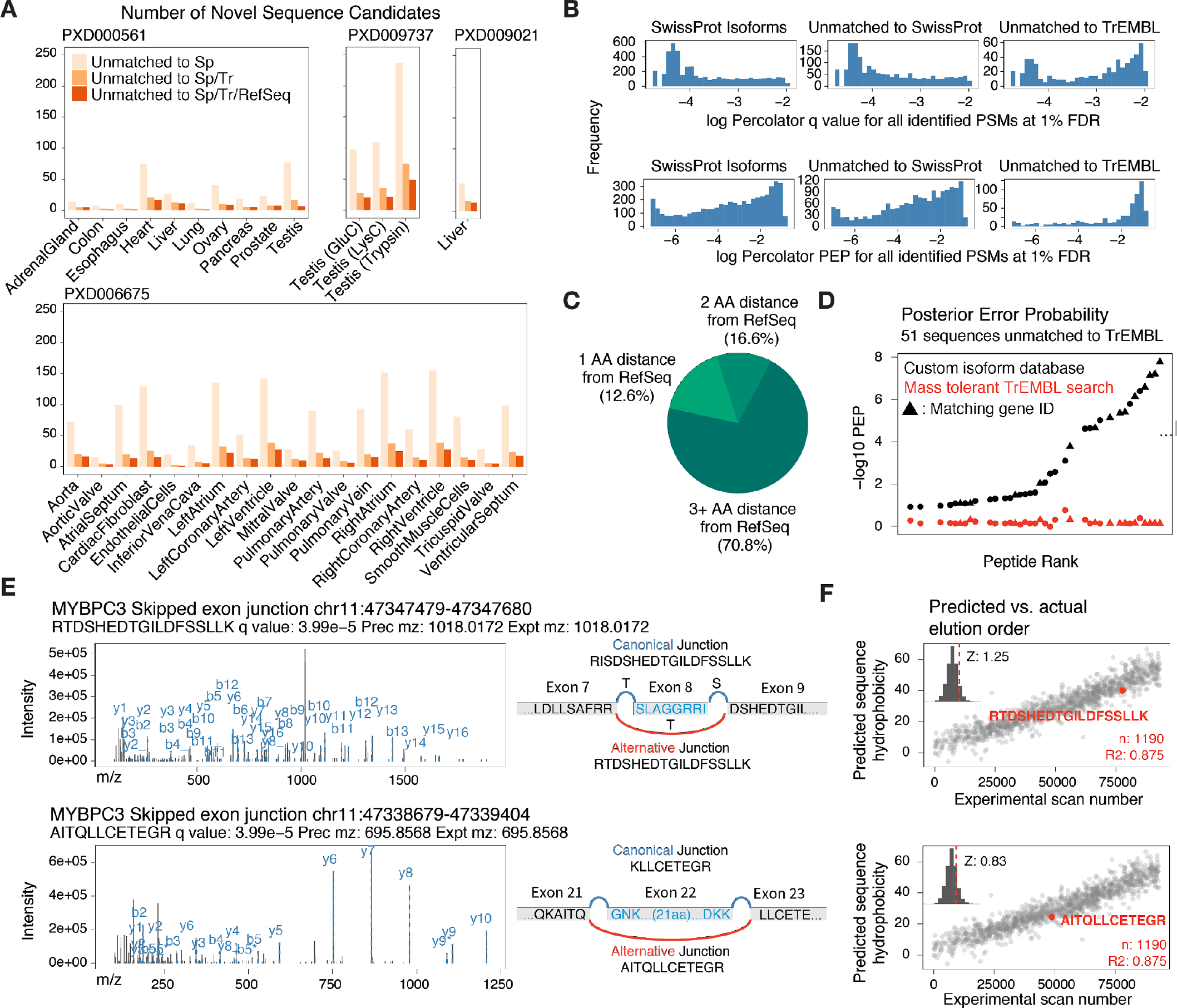
Splice junctions include novel peptides not documented in databases. **A.** Number of novel sequence candidates in each examined tissue. **B.** Proportion of novel sequences that are not mappable to RefSeq allowing 1, 2, or 3 mismatches. **C.** Comparison of −log10 Percolator posterior error probability for 51 TrEMBL unique sequence in the left ventricle against the results from the corresponding spectra in a mass tolerant open search using MSFragger against TrEMBL. **D.** Distribution of mass shifts and percolator FDR of non-canonical sequences that are matchable to SwissProt isoforms (left) against those that are not found in SwissProt (middle) or TrEMBL (right). **E.** Tandem mass spectra of two identified splice junction peptides (RTDSHEDTGILDFSSLLK and AITQLLCETEGR for MYPBC3) are shown. Neither peptide is matched to UniProt SwissProt/TrEMBL or Ensembl RefSeq sequences. **F.** Predicted hydrophobicity of the two novel sequences compared to high-quality spectra from the same experiment suggests the sequence eluted at the expected retention time when the spectrum was acquired. Inset: Z score of residuals from best-fit line.

First, we used a sequence alignment algorithm to assess whether the identified novel peptides may be matchable to RefSeq when allowing one or more mutations. We found that the majority (70.8%) of these sequences cannot be matched to a sequence in RefSeq even allowing 2 mismatches, suggesting the absolute majority of identified spectra are unlikely to arise from single amino acid variants differing from the reference proteome, or other unaccounted-for mass shifts at a single residue (Figure 4C). Second, we also evaluated whether the spectra may be better matched to a mass-tolerant open search for post-translational modifications (Figure 4D). We subjected the left ventricle dataset to a comprehensive MSFragger open search against TrEMBL, allowing −200 to +400 Da in mass shift, followed by Percolator filtering which identified 13,880 peptides from 8,702 protein groups at 1% FDR. Among the spectra identified to 51 novel peptide sequences in the left ventricle, 38 were matched to a sequence using MSFragger and 14 sequences were matched to the same gene ID, but only one spectrum were confidently identified at 1% FDR. The other spectra were not identified below FDR threshold (median Percolator q: 0.15) and all spectra had considerably higher adjusted P values than in the custom database search. By contrast, 267 out of the 394 (67%) non-canonical isoform sequences found in TrEMBL in the same sample were matched to identical sequences in the open search, suggesting the alternative splicing database search has additional identification power for a subset of spectra over modification-based open search.

Thirdly, we manually inspected tandem mass spectra of potentially novel junction peptides. Among the novel sequences are two skipped exons for myosin-binding protein C3 (RTDSHEDTGILDFSSLLK and AITQLLCETEGR), corresponding from skipping of exon 8 and exon 22, respectively (Figure 4E). Both sequences were identified from high-quality spectra with large proportions of matched fragment ions at Percolator q ≤ 0.01. We also determined whether the identified novel isoform peptides eluted in expected retention time based on their assigned sequences. Using an existing algorithm (SSRC) to determine the hydrophobicity coefficient of peptide sequences de novo (Krokhin et al., 2004), we benchmarked the novel peptides against high-confidence canonical peptides within the same experimental fraction. We prioritize novel peptides whose retention time residual to the best-fitted curve between predicted hydrophobicity and empirical retention time is within three standard standard deviations from the mean of that of high-quality peptides, as is the case for the two sequences shown here (Figure 4F). A subset of the novel sequence candidates appear to show unexpected retention time and are more likely to be false positives. We present the spectra and chromatograph for all candidate novel peptide sequences in **Supplementary Data 1**, and note that improving sequence elution time prediction algorithms may provide a useful determinant in adjudging the validity of sequence variants.

### Alternative protein isoforms overlap with disordered regions

We next inspected the potential proteome impact of the identified splice isoforms. Among the novel peptides we found a splice variant for myosin-binding protein C3 (MYBPC3) (Figure 4E). MYBPC3 is a 140 kDa cardiac protein which forms an important sarcomeric component in cardiomyocytes responsible for maintaining cardiac sarcomere structure, and which is one of the most commonly mutated genes in human familial hypertrophic cardiomyopathy. From a translated skipped exon event, we identified the splice junction peptide RTDSHEDTGILDFSSLLK which is not found in RefSeq, as well as its sister peptide TDSHEDTGILDFSSLLK. The sequences are identified widely in the tissues within the datasets analyzed here, including whole heart, left atrium, and left ventricle. UniProtKB/SwissProt catalogues two isoform protein entries for MYBPC3, including the SwissProt canonical entry with 1,274 residues and an alternative entry with 1,273 residues, in which the serine 408 and lysine 409 of the canonical entry are replaced by a single arginine. Neither entry encompasses the alternative isoform sequence we identified, which omits the segment of amino acid sequence SLAGGGRRIS from residues 275 to 284 encoded by exon 8, and which contains instead the identified peptide matching to residues 273 to 274 (RT-) of the canonical sequence, then resuming after the splice junction at residues 285 to 300 (-DSHEDTGILDFSSLLK). The noncanonical peptide has not been observed in the peptide repositories PeptideAtlas (containing 1.4 million distinct peptides) (Deutsch et al., 2015) or MassIVE-KB (2.3 million peptides) (Wang et al., 2018a), is not present in the common protein database UniprotKB SwissProt/TrEMBL, and has no identical match (100% sequence identity and coverage) to any sequences of any taxonomy in RefSeq via BLASTP (Boratyn et al., 2013).

Intriguingly, we found that this skipped exon falls within an unusual region of local disorder nested between but not overlapping two well-defined Ig-like protein domains. We asked whether the excised region overlapped with other structural features of interest, and found that the skipped exon falls on two of three successive phosphorylation sites (S275, S284, S304) that are known to be key regulatory sites in MYBPC3 by protein kinase A (PKA) (Figure 5A). The phosphorylation of S275, S284, and S304 in MYBPC3 by PKA as well as by other kinases is known to cause the MYBPC3 N-terminal domain to dissociate from myosin heavy chain and hence an increase in cardiac crossbridge formation (Rosas et al., 2015). Mutagenesis experiments in animal models replacing these serines with the phospho-negative mimetic alanine have found that the resulting hearts had abnormal relaxation velocity but not ejection fraction (Rosas et al., 2015), suggesting these sites exert diastolic regulations on the sarcomere and may be associated with diastolic dysfunction. Across the length protein sequence, the region excluded by the alternative junction is statistically enriched in known phosphorylation site (Fisher’s exact test P: 0.02).

**Figure 5.**
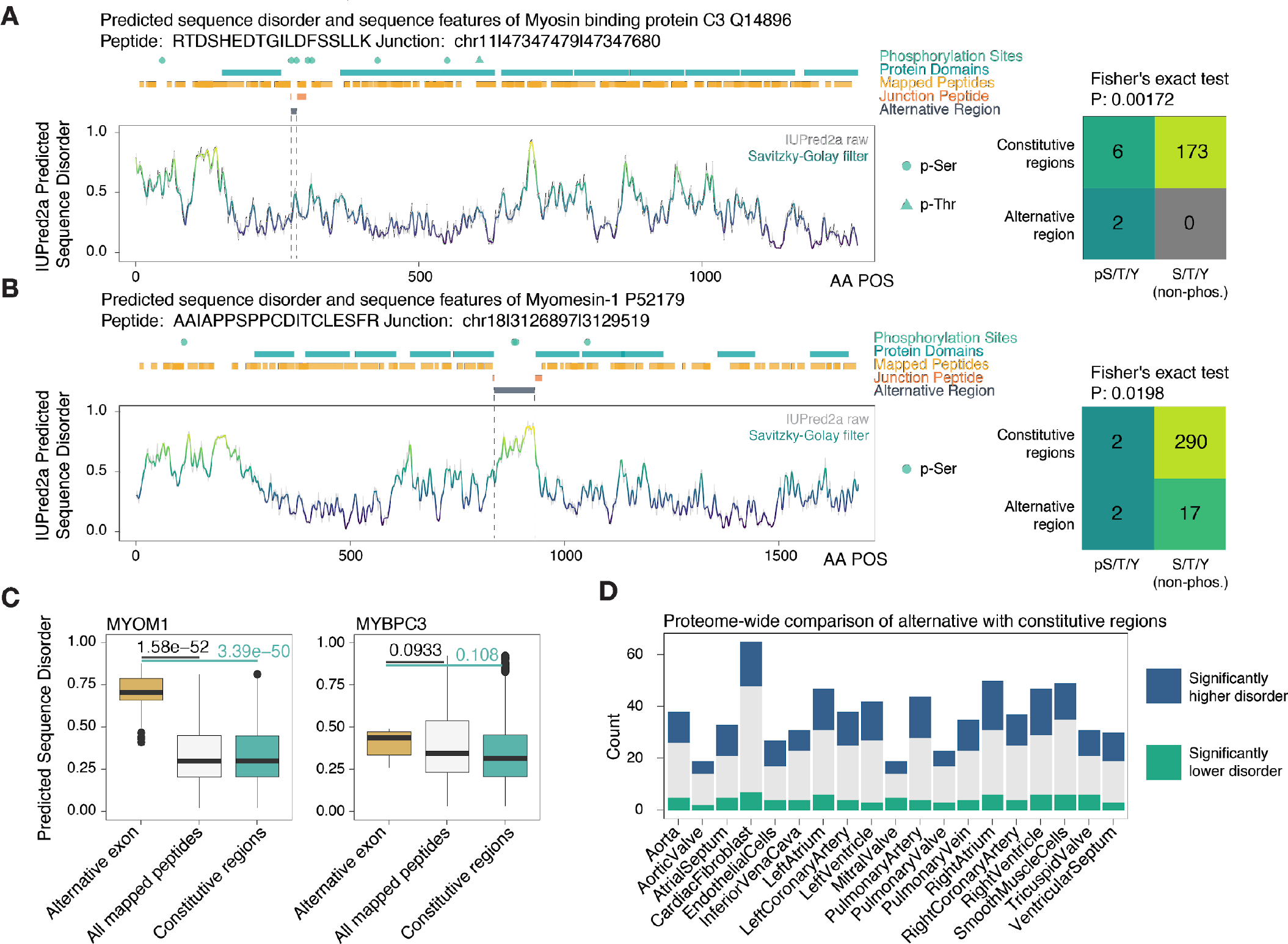
Splice isoforms preferentially overlap with disordered protein regions. **A.** Sequence features of myosin-binding protein C3 (MYBPC3), highlighting protein kinase A regulatory sites overlapping with the alternative region (residues skipped in the non-canonical isoform) of the protein, and the identified junction peptide spanning the excluded region. Sequence disorder was predicted de novo using IUPred2a and aligned with annotated protein domains and phosphorylation sites on UniProt. On the right, contingency table on the number of annotated phosphorylation sites and serine/threonine/tyrosine that are not annotated to be phosphorylated in the excluded region vs. the rest of the protein sequence. **B.** As above, for an alternative protein isoform of myomesin-1 (MYOM1). **C.** Box plots showing the distribution of sequence disorder in the alternative region (gold) of MYOM1 and MYBPC3 vs. all residues uniquely identified by peptide in the database search (white) and the full-length protein sequence excluding the alternative region (green). P value: Mann-Whitney test for differences in distribution. **D.** Over a proteome scale, there is an overrepresentation of alternative regions with significantly higher sequence disorder (blue) over the rest of the protein.

In another example where alternative isoforms overlap with important protein features, an isoform junction identified in myomesin-1 (MYOM1) with an excluded exon spanning a disordered region in between two fibronectin type III domains (Figure 5B). The alternative isoform differs from the rest of a protein with significantly higher sequence disorder (Mann-Whitney test for difference in distribution P: 3.3e-50) (Figure 5C). Other identified non-canonical sequences also showed a preferential location in high-disorder regions (**Supplementary Data 2**), such that on a proteome scale there is a clear preference for alternative isoforms to alter protein regions with heightened sequence disorder (Figure 5D). Taken together, the result provides evidence for one instance where protein alternative isoform that overlaps with known regulatory post-translational modification sites, and proteome-wide enrichment in disordered protein regions, suggesting two potential mechanisms through which alternative splicing may regulate proteome function.

### Non-canonical protein isoforms during cell differentiation

Lastly, we applied the workflow towards an in-house generated mass spectrometry dataset on human induced pluripotent stem cell (iPSC) differentiation into cardiomyocytes. Three human iPSC lines underwent directed cardiac differentiation over 14 days through an established small-molecule based protocol (Figure 6A). During the time course, we harvested cells daily and subjected cellular lysate and performed quantitative tandem mass spectrometry with the aid of 10-plex stable isotope-labeled tandem mass tags for MS2-based quantification (Supplementary Figure S1A-C). We then processed the mass spectrometry data files and identified cardiac-specific proteins using the human heart specific database generated above. The results reflected that cardiac protein expression profile correspond to the course of cellular differentiation and different stages of cardiac differentiation (Figure 6B). From 87 quantified non-canonical protein isoforms including 14 not in the SwissProt human canonical/isoform database, we observed that non-canonical isoforms exhibited diverse different stage-specific expression patterns (Figure 6C), including isoforms that are enriched in iPSCs, iPSC-derived-cardiomyocytes, and intermediary cell stages (Figure 6D). Comparing protein expression at multiple cell stages, we found cardiomyocyte specification is associated with altered expression of non-canonical isoforms particularly in actin binding and ribosomal processes (Figure 6E).

**Figure 6.**
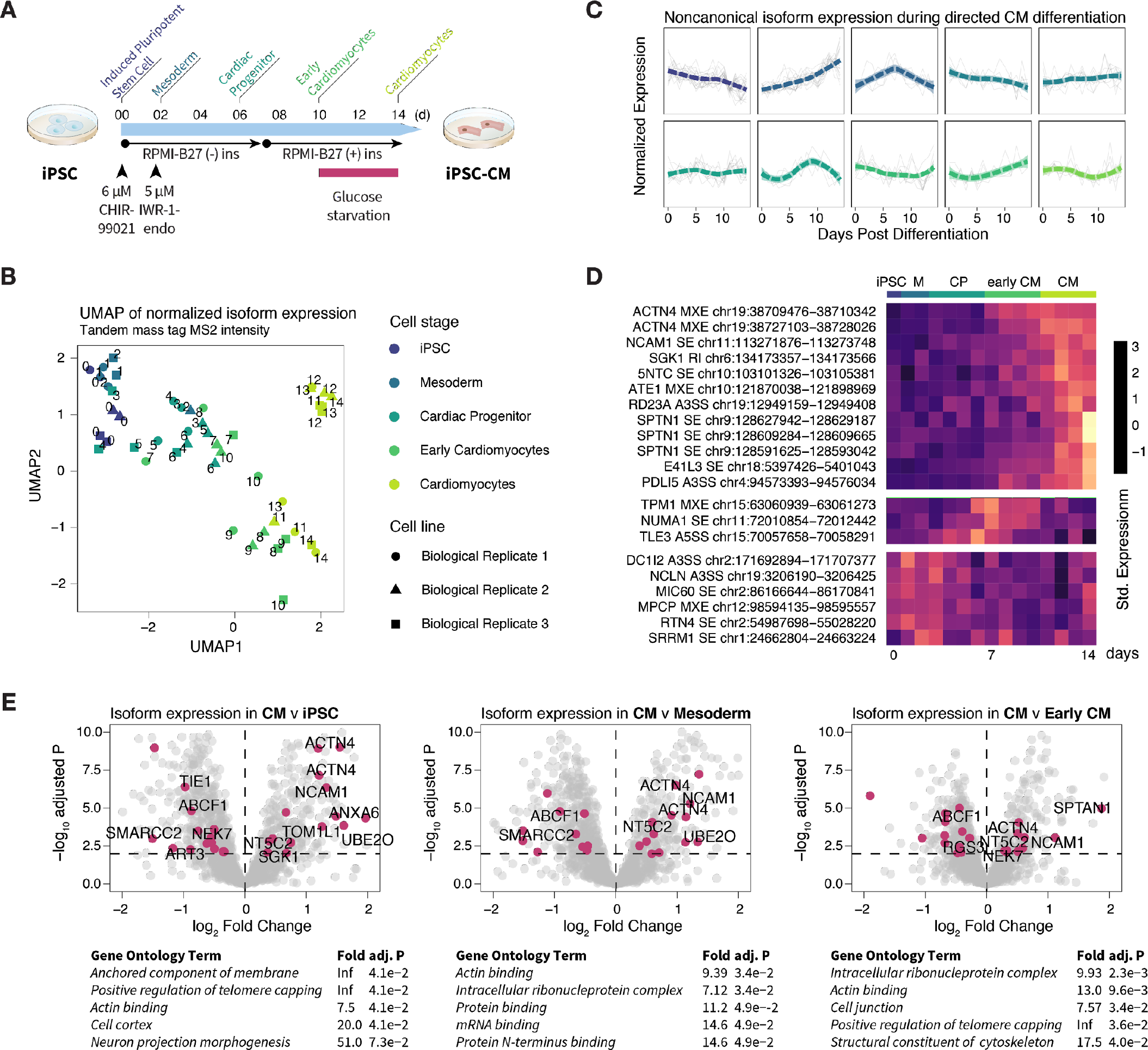
Expression of protein isoforms during directed cardiac differentiation. **A.** Schematic for human iPSC directed cardiac differentiation protocol with annotated stages (iPSC: day 0; mesoderm day 1-2; cardiac progenitor: day 3-6; early cardiomyocytes: day 7-10; cardiomyocytes: day 11-14). **B.** UMAP projection of tandem mass tag intensity shows that total protein expression reflects differentiation stages across three biological replicates. **C.** Hierarchical clustering of non-canonical peptide expression during cardiomyocyte differentiation shows diverse temporal behaviors of non-canonical isoforms in each cluster. **D.** Heat map showing the row-standardized expression of example non-canonical isoforms with cell-specific expression during differentiation. **E.** Volcano plot of log fold-change vs. −log adjusted P values showing comparison of protein expression between cardiomyocytes with (left) iPSC, (center) mesoderm, and (right) early cardiomyocytes. Data points: isoforms; magenta: differentially expressed non-canonical isoforms (limma adjusted P ≤ 0.01); differentially expressed isoforms not found in SwissProt are labeled.

To assess the effect of cell-type specific databases vs. the tissue-specific database in protein identification, we further generated deep RNA sequencing data containing approximately 100 million short-reads from day 0 and day 14 post-differentiation iPSCs and iPSC-derived cardiomyocytes. We used the resulting sequencing read to generate sample-specific protein sequence database using the method described here (Supplementary Figure S2A) We applied generated sample-specific protein sequence databases using RNA sequencing data from cells collected day 0 and day 14. The iPSC-specific databases show only partial overlap with human heart specific database suggesting the presence of cell type-specific translated junctions (Supplementary Figure S2B-C). Among genes with multiple quantified isoforms, the majority show concordant expression patterns during differentiation but we also observed isoform-specific changes (Supplementary Figure S2D-E). As an example, a skipped exon event in NDUFV3 created two quantified isoforms, whose expression differed in iPSC but converged during the course of differentiation. Altogether, the data suggest that non-canonical isoforms are dynamic to cell types and lineages, and our workflow may be applicable to extending profiling studies into cellular differentiation and other dynamic processes including in disease studies.

## Discussion

Alternative splicing is widely implicated in development of disease, but the protein molecular functions of most splice isoforms are uncharacterized. A clear picture on how alternative splicing impacts proteomes has yet to emerge but recent reports suggest that splicing can rewire interactomes (Yang et al., 2016) and also serve as a mechanism to decrease the abundance of the major canonical isoform of a gene (Liu et al., 2017). It is appreciated that only a minor subset of expressed transcripts have the potential to be translated, with about one-third of skipped exon events having been estimated to preserve protein structure and hence have higher translational potential (Hao et al., 2015), whereas the rest may be removed by nonsense-mediated decay or co-translational proteasomal degradation (Weatheritt et al., 2016). The ability to empirically discriminate which isoform transcripts exist at the protein level at a particular tissue will aid in elucidating the biochemical and signaling functions of alternative isoforms.

We describe here a splice-junction centric approach to create size-restricted databases to guide protein isoform identifications. The generation of accurate protein sequence databases is an important step in avoiding inflation of false positives during database search, and entails finding the set of isoform peptides that exists in a particular sample and is detectable by the mass spectrometry experimental design. A number of recent studies have demonstrated the use of RNA sequencing reads as a template to identify non-canonical protein sequences (Carlyle et al., 2018; Cifani et al., 2018; Zickmann and Renard, 2015), and proteogenomics analysis are increasingly common for large datasets that perform RNA sequencing and mass spectrometry in identical samples (Mertins et al., 2016; Wang et al., 2019). Our approach builds on prior work and is distinguished by the selection for splice junction pairs in alternative splicing events where there exists appreciable RNA read counts for both the exon-included and exon-excluded junctions. We also assume there is only one true translatable frame for each junction which is derived either from the known frame of the canonical form or choosing one frame that does not result in PTC during in silico translation. This resulted in size-restricted protein sequence databases that are a fraction of the sizes in typical approaches that utilize transcript assembly coupled to three-frame or six-frame translation (Ning and Nesvizhskii, 2010; Sheynkman et al., 2013; Wang and Zhang, 2013; Zickmann and Renard, 2015). In the RNA sequencing datasets analyzed here, the databases contained to 6.3% to 17.0% as many sequences as RefSeq. We demonstrated that the approach successfully recovered non-canonical isoforms across analyzed tissues including novel sequences not found in TrEMBL or RefSeq.

Many identified isoforms differ from their canonical counterpart in excluding regions that overlap sequence disorder and phosphorylation sites. A discovered myosin-binding protein C3 (MYBPC3) isoform differs from its canonical sequence by only in 10 of the 1,274 residues, but is located at a crucial phosphorylation region known to modulate diastolic functions of the heart, which suggests a potential manner through which it can impact protein function. It has been observed that the majority of alternative exons do not alter conserved stable protein domains (Barbosa-Morais et al., 2012; Buljan et al., 2012), which has been cited as evidence against the importance of alternative splicing to the proteome (Tress et al., 2017b, 2017a). However, unfolded protein regions also play important roles in regulating protein functions including by forming protein-protein interaction surfaces (Ellis et al., 2012), governing phase separation and membraneless organelles (Harmon et al., 2017; Uversky, 2017), and exerting allosteric modulation on remote catalytic domains (Ferreon et al., 2013; Keul et al., 2018). Taken together, our results support the notions that on a proteome scale splicing can influence proteome function by (i) rewiring flexible regions linking stable protein domains; and (ii) provide a separate PTM control mechanism by toggling the binary presence/absence of phosphorylatable sites. The intersection between alternative splicing and sequence disorder and PTMs has been theorized (Zhou et al., 2018) and is consistent with prior suggestions that alternative isoforms alter protein interaction landscapes (Ellis et al., 2012; Yang et al., 2016).

In summary, we describe an approach to create more concise splice variants databases for protein isoform analysis by considering alternative splicing events in a sample. Among uniquely mappable distinct peptide sequences, we found that 1 to 3% mapped to non-canonical isoforms per tissue. Additional translated splice junctions likely remain to be discovered. A proportion of alternative junction peptides appear in multiple custom translated forms due to the combinatorial redundancy in exon junctions, rendering them ambiguous in protein assignment. Increasing performance and adoption of long-read RNA sequencing and middle-down/top-down proteomics will help mitigate this limitation. Finally, continued refinements in computational methods to predict translated transcripts will contribute to ongoing efforts in protein isoform identification.

## Methods

### Public RNA sequencing and mass spectrometry datasets

RNA sequencing datasets were retrieved from ENCODE at the following accessions: heart (ENCSR436QDU, ENCSR391VGU), liver (ENCSR226KML, ENCSR504QMK), lung (ENCSR425RGZ, ENCSR406SAW), pancreas (ENCSR671IYC, ENCSR586SYA), adrenal gland (ENCSR801MKV, ENCSR754WLW), transverse colon (ENCSR800WIY, ENCSR403SZN), ovary (ENCSR841ADZ, ENCSR042GYH), esophagus (ENCSR098BUF, ENCSR750ETS), testis (ENCSR029KNZ, ENCSR344MQK), and prostate (ENCSR495HDM, ENCSR701TST). RNA sequencing data from at least two biological replicates from each tissue were used. All data were 101nt paired-end total RNA sequencing generated on an Illumina Hi-Seq 2500 sequencer and passed ENCODE quality control (ENCODE Project Consortium, 2012). RNA sequencing read [.fastq] files were manually retrieved on 2017-11-12. Proteomic datasets in Thermo [.raw] format were retrieved from ProteomeXchange/PRIDE (Deutsch et al., 2017) at the following accessions: “A draft map of the human proteome” (PXD000561) (Kim et al., 2014) generated on Thermo Orbitrap Velos and Orbitrap Elite mass spectrometers with FT/FT; “Region and cell-type resolved quantitative proteomic map of the human heart and its application to atrial fibrillation” (PXD006675) (Doll et al., 2017) generated on a Thermo Q-Exactive HF mass spectrometer; “Human testis off-line LC-MS/MS” (PXD009737) (Sun et al., 2018) generated on a Thermo Q-Exactive HF-X mass spectrometer; and “Proteomic analysis of human liver reference material” (PXD009021) generated on a Thermo Fusion Lumos mass spectrometer in FT/FT mode.

### RNA data processing and database generation

To align the retrieved RNA sequencing data, we used STAR v.2.5.0a (Ballouz et al., 2018; Dobin and Gingeras, 2016) on a Linux 4.10.0-32-generic Ubuntu x86_64 workstation. We mapped .fastq sequences to Ensembl GRCh38.89 STAR indexed genomes with Ensembl .gtf annotations (--sjdbGTFfile GRCh38.89.gtf --sjdbOverhang 100). To extract splice junctions from the mapped and compare splice levels across biological replicates, we used rMATS-Turbo v.0.1 (Shen et al., 2014) on the mapped bam files with the following options (--readLength 101 --anchorLength 1). We implemented a custom script written in-house in Python v.3.6.1, which tabulates the rMATS results on alternative splicing events from each tissue and filters out ineligible splice pairs by virtue of read count threshold or significant inter-sample differences. Junctions are filtered by the minimal excluded junction read count of all biological replicates (rMATS *SJC*) for a particular junction *j* for a tissue *t* such that transcript level *SJC*_*j,t*_ is above a detectability threshold *SJC*_*j,t*_ ≥ *θ*_*t*_, which is estimated by a mixture Gaussian model of excluded junction read counts based on the specific RNA sequencing dataset for the tissue. In addition we assume that the isoform is reliably observed across multiple runs, employing the statistical model implemented in rMATS and exclude significantly differential splice junctions at P ≤ 0.01 (Shen et al., 2014).

The script next retrieves nucleotide sequences from each splice pair based on the recorded genomic coordinates using the Ensembl REST web application programming interface, and attempts to identify the appropriate translation frames, transcription start sites, and transcription end sites of each splice pair from the Ensembl GRCh38.89 annotation GTF file based on the upstream exon. The retrieved qualifying nucleotide sequences are further translated into amino acid sequences using the annotated phase and frame. Peptides are selected for inclusion if they fulfill one of the following sequential considerations: (i) they are translated in-frame by the GTF-annotated translation frame in Ensembl GRCh38.89 GTF successfully without encountering a frameshift or PTC; or (ii) one of the spliced pair junctions encountered a frameshift event using the GTF-annotated frame but both are translated without PTC; (iii) they are translated without PTC using a single translation frame different from the GTF-annotated frame; (iv) in rare occasions, one of the two junctions but not the other encountered a PTC. Finally, all translated peptides are required to be stitchable back to the SwissProt canonical sequences retrieved via the gene name using a 10-amino-acid overhang. Orphan peptides that are translated but not stitchable back to SwissProt are discarded from the analysis. The translated databases used for analysis are available in **Supplementary Data 3**.

### Mass spectrometry database search and analysis

Mass spectrometry raw spectrum files were converted to open-source [.mzML] formats using ProteoWizard msconvert v.3.0.11392 (Adusumilli and Mallick, 2017) with the following options (--filter “peakPicking vendor”). Database search against custom databases were performed using the SEQUEST algorithm implemented in Comet v.2017.01 rev.0 (Eng et al., 2015) with the following options (--peptide_mass_tolerance 10 --peptide_mass_unit 2 --isotope_error 2 --allowed_missed_cleavage 2 --num_enzyme_termini 1 --fragment_bin_tol 0.02). Conventional settings for other Comet parameters were used and a reverse decoy database was generated from the custom database for each search for FDR estimation. Static cysteine carboxyamidomethylation (C +57.021464Da; Unimod accession #4) modification was specified. Tryptic and semi-tryptic peptides within a 10-ppm parent mass window surrounding the candidate precursor mass were searched, allowing up to 2 miscleavage events.

Peptide spectrum match data were filtered and target and decoy sequence matches were re-ranked using the semi-supervised learning method implemented in Percolator (The et al., 2016) in the Crux v.3.0 Macintosh binary distribution (McIlwain et al., 2014) with the following options (--protein T --fido-empirical-protein-q T --decoy-prefix DECOY_). Peptides with Percolator q value ≤ 0.01 are considered to be confidently identified. Mass tolerant open search comparison was performed using MSFragger (Kong et al., 2017) using standard parameters with lower mass tolerance −200 Da and upper mass tolerance was +400 Da against the UniProt TrEMBL human database (accessed 2019-02-08), followed by Percolator filtering as above.

### Human iPSC culturing and RNA sequencing

Human iPSCs were acquired from cryopreserved stocks in the Stanford Cardiovascular Institute Biobank and expanded in monolayer in Gibco Essential 8 medium (Thermo) on a Matrigel matrix (Corning). Directed differentiation into cardiomyocytes was performed on three individual donor lines using an established small-molecule Wnt-activation/inhibition protocol yielding 95% pure TNNT2+ cardiomyocytes (Burridge et al., 2014). Briefly, iPSC cultures at ~90% confluence in 6-well-plates were treated with 6 μM CHIR-99021 (SelleckChem) in RPMI 1640 medium supplemented with B27 supplements (Thermo Fisher Scientific) for 2 days to induce mesoderm specification, allowed to recover 1 day, then treated with 5 μM IWR-1-endo (SelleckChem) for 2 days for cardiac specification. On day 7, the culture medium was changed to RPMI-B27 + insulin, and the cardiomyocytes were glucose-starved on day 10 to day 14. Cells were harvested daily at day 0 to day 14 post-differentiation by dissociation using TrypLE select 10x (Thermo Fisher Scientific) and pelleted by centrifugation (200 ×g, ambient temperature, 5 min).

Total cellular RNA from day 0 and day 14 post-differentiation were extracted by 300 μL TRIzol/chloroform per ~1e6 cells, followed by solid-phase extraction using RNeasy mini columns (QIAgen) according to the manufacturer’s protocol. Purified RNA was eluted in 50 μL of RNAse-free water and the yield quantity and quality were assessed by fragment electrophoresis on an Agilent Bioanalyzer with the RNA Integrity Number (RIN) of all samples used for sequencing above 9.0. RNA sequencing was performed on an Illumina Hi-Seq instrument to acquire paired-end 150-nt reads up to a read-depth of 31.1G to 41.7G clean bases (Novogene). The RNA sequencing data were processed identically to the public datasets above to create a custom FASTA database containing the combined human alternative splice junctions from both day 0 and day 14 time points.

### Tandem mass tag labeled quantitative mass spectrometry

Cell lysate proteins from each iPSC time point were extracted by commercial RIPA or M-Per tissue lysis buffer (Thermo Fisher Scientific) followed by brief pulses of sonication with typically 6 pulses at 20% amplitude followed by 5 sec cooldown on ice. Total protein extracts for each sample were quantified by bicinchoninic acid assays and 100 μg proteins were digested on 10-kDa MWCO polyethersulfone filters (Thermo Fisher Scientific). Samples were washed with 8 M urea, buffer-exchanged with triethylammonium bicarbonate (50 mM, 500 μL), reduced with tris(2-carboxyethyl)phosphine (10 mM, 70°C, 5 min) and alkylated with iodoacetamide (18mM, ambient temperature, 30 min). Proteins were digested on-filter (16 hr, 37 °C) with sequencing-grade modified trypsin (50:1, Promega). Proteolysis was terminated and peptides eluted with 20 μL of 10% trifluoroacetic acid followed by centrifugation (13,000 ×g, ambient temperature, 15 min). Digests were quantified using a colorimetric peptide quantification kit (Thermo Fisher Scientific) and labeled with 10-plex tandem mass tags (Thermo Fisher Scientific) at ambient temperature with 600 rpm shaking for 2 hr. Label assignment was randomized using a random number generator. Labeling was quenched with 5% hydroxylamine.

Liquid chromatography-tandem mass spectrometry was performed on peptides fractionated into 6 fractions using pH-10 reversed-phase spin columns (Thermo Pierce). Second-dimension liquid chromatography was performed using a single Easy-nLC 1000 nanoflow ultrahigh-pressure liquid chromatography (UPLC) system on an EasySpray C18 column (PepMap, 3-μm particle, 100-Å pore; 75 μm × 150 mm; Thermo Fisher Scientific) in 120-min in a pH-2 reversed-phase gradient. The nano-UPLC was run at 300 nL/min with the gradient of 0 to 105 min, 0 to 40 %B, 105 to 110 min, 40 to 70 %B, 110 to 115 min, 70 to 100 %B, hold for 5 min, with solvent B being 80% v/v acetonitrile and 0.1% v/v formic acid. Mass spectrometry was performed using a Q-Exactive HF high-resolution Orbitrap mass spectrometer (Thermo Fisher Scientific) coupled to the nano-UPLC by an EasySpray interface. Each MS1 survey scan was acquired at 120,000 resolving power in positive polarity in profile mode from 350 to 1650 m/z, lock mass, dynamic exclusion of 90 sec, maximum injection time of 20 msec, and automatic gain control target of 3e6. MS2 scans were acquired on the top 15 ions with monoisotopic peak selection at 60,000 resolution, automatic gain control target of 2e5, maximum injection time of 110 ms, and isolation window of 1.4 m/z, with typical normalized collision-induced dissociation energy of 35 or stepped normalized collision-induced dissociation energy of 27, 30, and 32.

### Additional data analysis and statistics

To quantify peptide intensity in the iPSC data, tandem mass tag intensity was corrected by the isotope contamination matrix supplied by the manufacturer, tag intensity in each 10-plex experiment was column normalized, row-normalized by two pooled reference tags per experimental block, then normalized by trimmed means of m values in edgeR (Robinson and Oshlack, 2010) and log-transformed for across-sample comparison. Non-unique peptides as well as peptides confidently identified at fewer than three independent tandem mass tag experiment blocks were discarded. Statistical analysis of differential expression was performed using the regularized t-test and empirical Bayes model in limma (v.3.34.3) in R/Bioconductor (v.3.6) (Ritchie et al., 2015) using discrete developmental stages as factors. Proteins with limma adjusted P value (FDR) ≤ 0.01 in each comparison are considered to show evidence for statistically significant differential regulation.

Data statistical analysis and visualization were performed in R v.3.4.4 (2018-03-15 release) or above on x86_64-apple-darwin15.6.0 (64-bit) with the aid of Bioconductor v.3.6 (Huber et al., 2015), and MSnBase v.2.4.2 (Gatto and Lilley, 2012). Gene Ontology terms were used for protein functional annotations (The Gene Ontology Consortium, 2017). Protein sequence features were retrieved from UniProt (UniProt Consortium, 2018). Protein sequence disorder prediction was performed using IUPred2A (Mészáros et al., 2018). Fisher’s exact test was used to assess enrichment in phosphorylation sites in isoform excluded regions and in the enrichment of Gene Ontology terms in quantified proteins. Sequence occurrence of identified peptide sequences in UniProt SwissProt or TrEMBL human (9606) sequences (retrieved 2019-02-08) (UniProt Consortium, 2018) or Ensembl RefSeq (retrieved 2019-02-07) (Pruitt et al., 2014) with 0 or more mismatch tolerance were assessed using the BioStrings v.3.7.0 package. Dimension reduction of human iPSC tandem mass tag data was performed using the uniform manifold approximation and projection (UMAP) method as described (Becht et al., 2018).

## Acronyms and Abbreviations

A3SS: alternative 3-prime splice site
A5SS: alternative 5-prime splice site
FDR: false discovery rate
IDR: intrinsically disordered regions
iPSC: induced pluripotent stem cells
MXE: mutually exclusive exons
PSI: percent spliced in
PTC: premature termination codon
PTM: post-translational modifications
SE: skipped exon
RI: retained intron
TMT: tandem mass tags

## Acknowledgments

This work was supported in part by National Institutes of Health (NIH) research grants F32 HL139045 and K99 HL144829 to E.L.; T32 HL007822 (CU AMC) for Y.H.; R01 HL141371 and R01 HL146690 to J.C.W.; R01 GM117624, R01 HL141278, and The University of Colorado Consortium for Fibrosis Research and Translation Pilot Grant to M.P.L.

## Supporting Information

### Supplementary Data 1

Spectral and chromatographic evidence of identified novel peptides. Figshare https://doi.org/10.6084/m9.figshare.6493247.v2

### Supplementary Data 2

Protein sequence features along alternative splice junctions. Figshare https://doi.org/10.6084/m9.figshare.6493262.v2

### Supplementary Data 3

Human tissue-specific protein sequence FASTA database. Figshare https://doi.org/10.6084/m9.figshare.7780940.v1

---

**Supplementary Figure S1.**
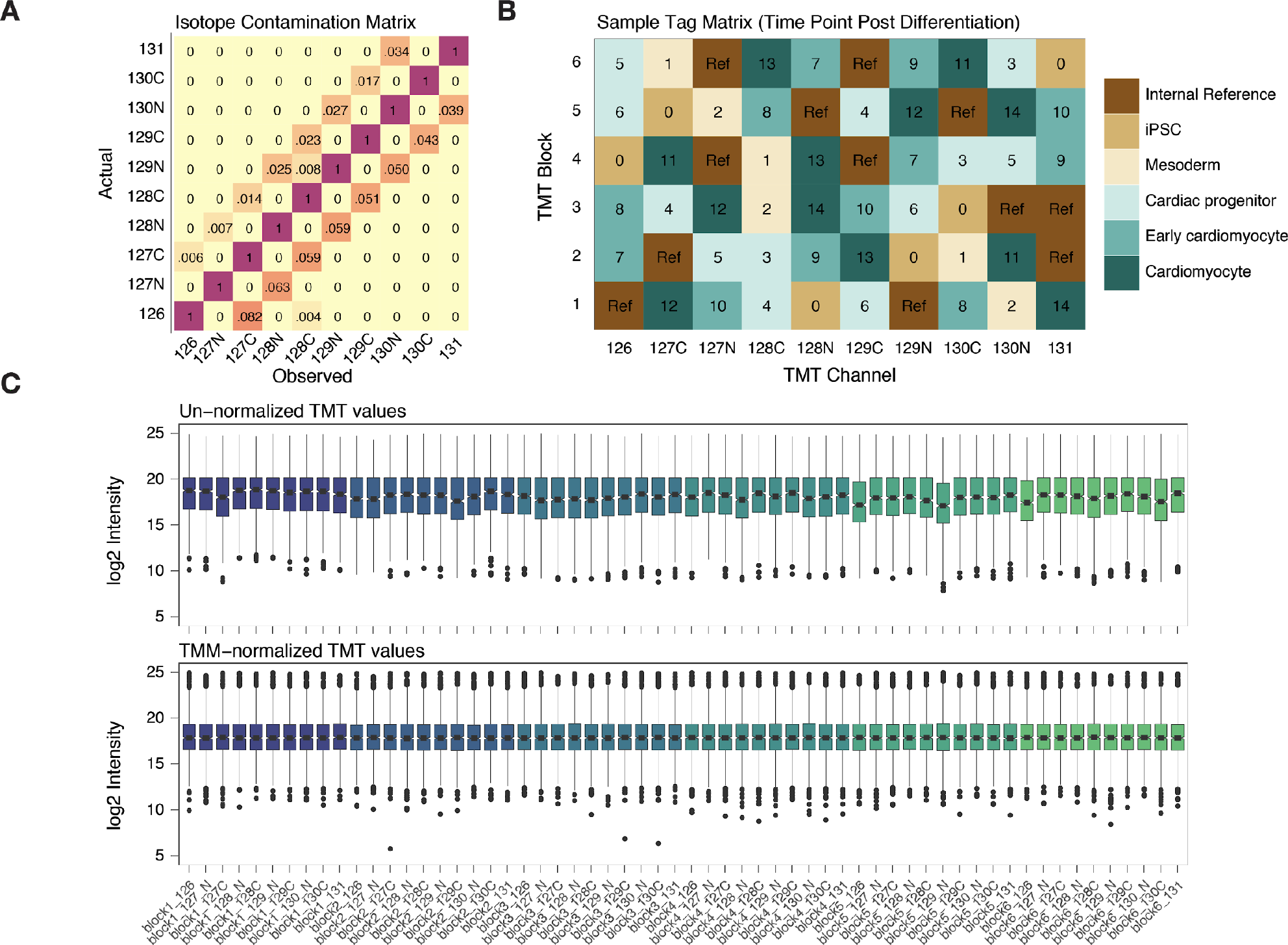
Tandem mass tag labeling of human iPSC proteins. **A.** Isotope cross-contamination matrix for calculating actual channel intensity from observed tag intensity in tandem mass tag (TMT) data. **B.** Randomized sample assignment and internal references for TMT channels. Labels denote time point (days) post differentiation or pooled internal reference (Ref). Fill color denotes discretized differentiation stages. **C.** Distribution of unnormalized (upper) and column- and trimmed mean of M value (TMM)- normalized (lower) TMT intensity for each channel in each experimental block used for differential expression analysis.

**Supplementary Figure S2.**
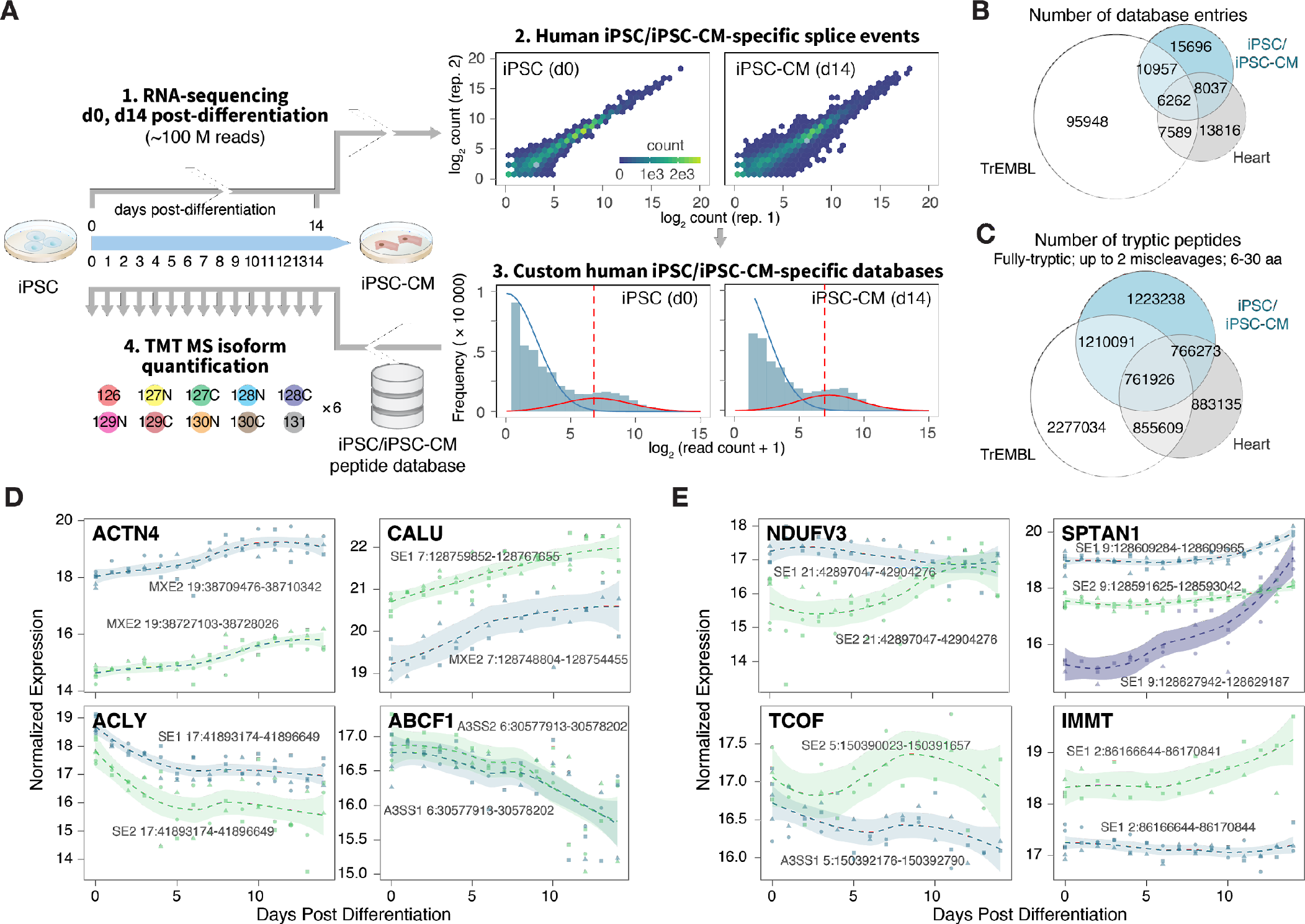
Human iPSC specific isoform sequence database. **A.** Experimental schema. Deep RNA sequencing data were generated from day 0 and day 14 iPSC and iPSC-cardiomyocytes, respectively. Cell specific databases are used to re-process the time-course tandem mass tag data. **B-C** The cell-specific databases show partial overlaps with the human heart-specific databases in **B.** database entries and **C.** tryptic peptides (6 to 30 amino acids, allowing one miscleavages). **D-E.** Isoforms from the same gene may show **D.** concordant or **E.** discordant expression patterns during cardiomyocyte differentiation. Trendline and shaded areas show local regression (loess) and bootstrap uncertainty regions.

